# FOD-Net: A Deep Learning Method for Fiber Orientation Distribution Angular Super Resolution

**DOI:** 10.1101/2021.01.17.427042

**Authors:** Rui Zeng, Jinglei Lv, He Wang, Luping Zhou, Michael Barnett, Fernando Calamante, Chenyu Wang

## Abstract

Mapping the human connectome using fiber-tracking permits the study of brain connectivity and yields new insights into neuroscience. However, reliable connectome reconstruction using diffusion magnetic resonance imaging (dMRI) data acquired by widely available clinical protocols remains challenging, thus limiting the connectome / tractography clinical applications. Here we develop fiber orientation distribution (FOD) network (FOD-Net), a deep-learning-based framework for FOD angular super-resolution. Our method enhances the angular resolution of FOD images computed from common clinical-quality dMRI data, to obtain FODs with quality comparable to those produced from advanced research scanners. Super-resolved FOD images enable superior tractography and structural connectome reconstruction from clinical protocols. The method was trained and tested with high-quality data from the Human Connectome Project (HCP) and further validated with a local clinical 3.0T scanner. Using this method, we improve the angular resolution of FOD images acquired with typical single-shell low-angular-resolution dMRI data (e.g., 32 directions, *b* = 1000 s/mm^2^) to approximate the quality of FODs derived from time-consuming, multi-shell high-angular-resolution dMRI research protocols. We also demonstrate tractography improvement, removing spurious connections and bridging missing connections. We further demonstrate that connectomes reconstructed by super-resolved FODs achieve comparable results to those obtained with more advanced dMRI acquisition protocols, on both HCP and clinical 3.0T data. Advances in deep-learning approaches used in FOD-Net facilitate the generation of high quality tractography / connectome analysis from existing clinical MRI environments. Our code is freely available at https://github.com/ruizengalways/FOD-Net.

## 1. Introduction

The human brain connectome, which maps structural brain connectivity *in-vivo* through non-invasive diffusion MRI (dMRI) techniques, yields invaluable insights for the frontier research fields of brain-inspired artificial intelligence, psychiatry, neurology, and neurosurgery Savage (2019); Tozzi et al. (2020); Craddock et al. (2013). The reliability of dMRI-based connectomics is heavily dependent on the diffusion model employed for estimating local fiber orientations. The spherical deconvolution model Tournier et al. (2004, 2007); Jeurissen et al. (2014), with its inferred fiber orientation distribution (FOD) data, can be used to disentangle complex fiber structures within a voxel, and is currently considered among the state-of-the-art for tractography applications Dell’Acqua and Tournier (2019). It is optimally applied to multi-shell (i.e. data acquired at multiple *b*-values), high angular resolution diffusion imaging (HARDI, i.e. dMRI data acquired with a large number of diffusion gradient directions at each *b*-value, e.g. typically over 60 directions) data. However, the availability of multi-shell HARDI data in clinical settings is hampered by scanner and protocol limitations and practical constraints on total acquisition time, thus limiting broad application in both clinical and clinical research environments. In this regard, single-shell low angular resolution diffusion imaging (LARDI) data are more commonly used in clinical settings, and often combined with lower *b*-values (e.g. *b* = 1000 s/mm^2^). However, LARDI acquisitions cannot reliably resolve complex white matter fiber bundles configurations due to insufficient angular resolution, producing both spurious and missing connections Farquharson et al. (2013). This can degrade the connectome reconstruction, potentially impacting the validity of conclusions drawn from such data.

A growing interest in healthy and diseased brain connectomics has highlighted a critical need to bridge the gap between high-angular resolution, multi-shell dMRI acquisitions and the restricted practical capacity of clinical MRI investigations. Pioneering work Scherrer et al. (2011, 2012); Lin et al. (2019); Tian et al. (2020) has attempted to overcome the limitations of clinical protocols by striving for dMRI reconstruction, which in turn reconstruct more reliable fiber tracking connectivity information from dMRI itself. However, these methods require specific dMRI acquisition protocols (for example, a certain number of gradient directions), which is inflexible and impractical in general clinical environments. Specifically, since the input size of their deep learning models Lin et al. (2019); Tian et al. (2020) is fixed, they cannot cope with dMRI images with different number of gradient directions in inference stage, and as such that these methods have limited applicability given the variance of dMRI data size reflecting a diversity of acquisitions protocols. Moreover, given the nature of dMRI data possessing high inter-scanner and intra-scanner variance, performing reconstruction using raw dMRI images would suffer from the domain shift problem Ghafoorian et al. (2017) and as such that the performance of models is prone to be degraded. Here we present fiber orientation distribution (FOD) network (FOD-Net), a deep-learning-based FOD angular super resolution method that directly enhances FOD data from singleshell LARDI computed from typical clinical-quality data, to obtain the super-resolved FOD data equivalent to those derived from high-quality multi-shell (MS) HARDI acquisitions, thereby enabling reliable structural connectome reconstruction using widely available clinical protocols. Our core insight is that we can reformulate this research goal as the FOD angular super-resolution task rather than the dMRI angular superresolution task as such that we can bypass the direct use of dMRI data. Specifically, FOD data, which is taken as input by FOD-Net, is represented by the coefficients of spherical harmonic expansion Tournier et al. (2004). FOD data can be set to have the same data matrix size regardless of the number of diffusion directions used in the dMRI acquisition protocol (e.g. by setting the maximum harmonic order, *l_max_*, to a common value). Furthermore, in contrast to existing works establishing relationship between two completely different domains, i.e., dMRI domain and FOD domain, the proposed formulation aims to enhance and refine single-shell LARDI FOD images within the FOD domain itself. Moreover, thanks to multi-tissue informed FOD intensity normalisation Raffelt et al. (2017), quantitative FOD comparison across different subjects is enabled and therefore the domain shift problem caused by the direct use of raw dMRI is significantly alleviated. Our contributions are summarized as follows:

1. To our best knowledge, this is the first work which defines the FOD angular super-resolution task to generate super-resolved FOD images from low-quality single-shell LARDI FOD data, which is widely available in clinical environments. In contrast to dMRI reconstruction formulation, the FOD angular super-resolution task avoids directly using dMRI data as such that it has significant flexibility in terms of inter-scanner and intra-scanner variance. The preprocessing steps and performance measurement criteria for fiber orientation distribution angular super resolution have been established to facilitate the prosperity of future research community.
2. We proposed FOD-Net for fiber orientation distribution angular super-resolution task. It naturally takes as input a given single-shell LARDI FOD image and output the super-resolved FOD image. Super-resolved FOD images remove spurious fibers and recovers missing fiber tracks existing in the original single-shell LARDI FOD images and therefore enables reliable tractography and structural connectome reconstruction in practical clinical settings. FOD-Net
3. Extensive experiments have been conducted in the challenging datasets including the Human Connectome Project (HCP) and the single-shell LARDI dataset acquired at a local clinical 3.0T scanner to demonstrate the performance of FOD-Net. Compared to the state-of-the-art single-sell 3-tissue (SS3T) constrained spherical deconvolution (CSD) method Dhollander et al. (2016); Khan et al. (2020), we demonstrate the success of FOD-Net by improving the accuracy of each step in the connectome reconstruction pipeline, including FOD angular superresolution (see Fig. 2 for illustration), tractography, and connectome generation.

## 2. Materials and methods

### 2.1. Problem formulation

Suppose that we are given a single-shell (SS) LAR dMRI image 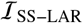 of size *W × H × L × N*, where *W, H, L*, and *N* are the width, the height, the length, and the number of gradient directions, respectively. Subsequently, SS3T CSD is used to compute the corresponding FOD image 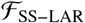, which is of size *W × H × L × C*, from 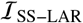, where *C* is the number of coefficients used to represent a FOD using the spherical harmonic basis. *C* can be set to have the same size regardless of the number of diffusion directions used in the dMRI acquisition protocol (e.g. by setting the maximum harmonic order, *l_max_*, to a common value to determine *C*). The task of fiber orientation distribution angular super-resolution is to enhance 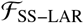 to obtain the super-resolved version 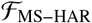, which is supposed to be generated from multi-shell (MS) HARDI dMRI image 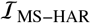 using multi-shell multi-tissue (MSMT) constrained spherical deconvolution (CSD) Jeurissen et al. (2014). 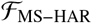 is supposed to have the same size as 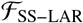, i.e., 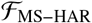 is of size *W × H × L × C*, while 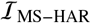 may has different size *W × H × L × M* from 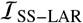 due to the variance of dMRI acquisition. Here 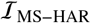 is supposed to be acquired using much more sophisticated dMRI protocols (typically multi-shell HARDI protocols) than 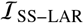 on the same subject, and as such that *M* is much greater than *N*.

Instead of reconstructing 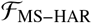 using 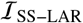, we reformulate the fiber orientation distribution angular superresolution task, which aims to find a function *f_θ_* with parameters *θ* that maps every single voxel of 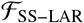 to its corresponding super-resolved voxel of 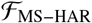, leading to:

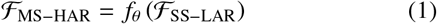

*θ* can be obtained by minimizing the following object function:

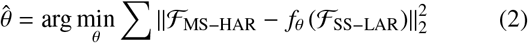

This problem is ill-posed because each 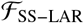 may correspond to multiple 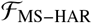 due to noise patterns introduced by different scanners. In this study, we parameterise the function *f* as the proposed fiber orientation distribution network (FOD-Net) (see Section 2.2 for details), which aims to tackle this challenging FOD angular super-resolution task in an end-to-end way.

### 2.2. FOD-Net architecture

Fig. 1 shows the schematic view of our FOD-Net in a clinical connectome reconstruction pipeline and its network architecture. Given a single-shell LARDI dMRI image acquired using a clinical protocol, SS3T CSD is first used to compute the singleshell LARDI FOD image. Then FOD-Net is applied to the computed single-shell LARDI FOD image to obtain the corresponding super-resolved FOD image with quality comparable to one produced from advanced research scanners. Subsequently, the tractography and connectome reconstruction are performed on the super-resolved FOD image to obtain the reliable tractogram and the connectome matrix.

**Fig. 1.**
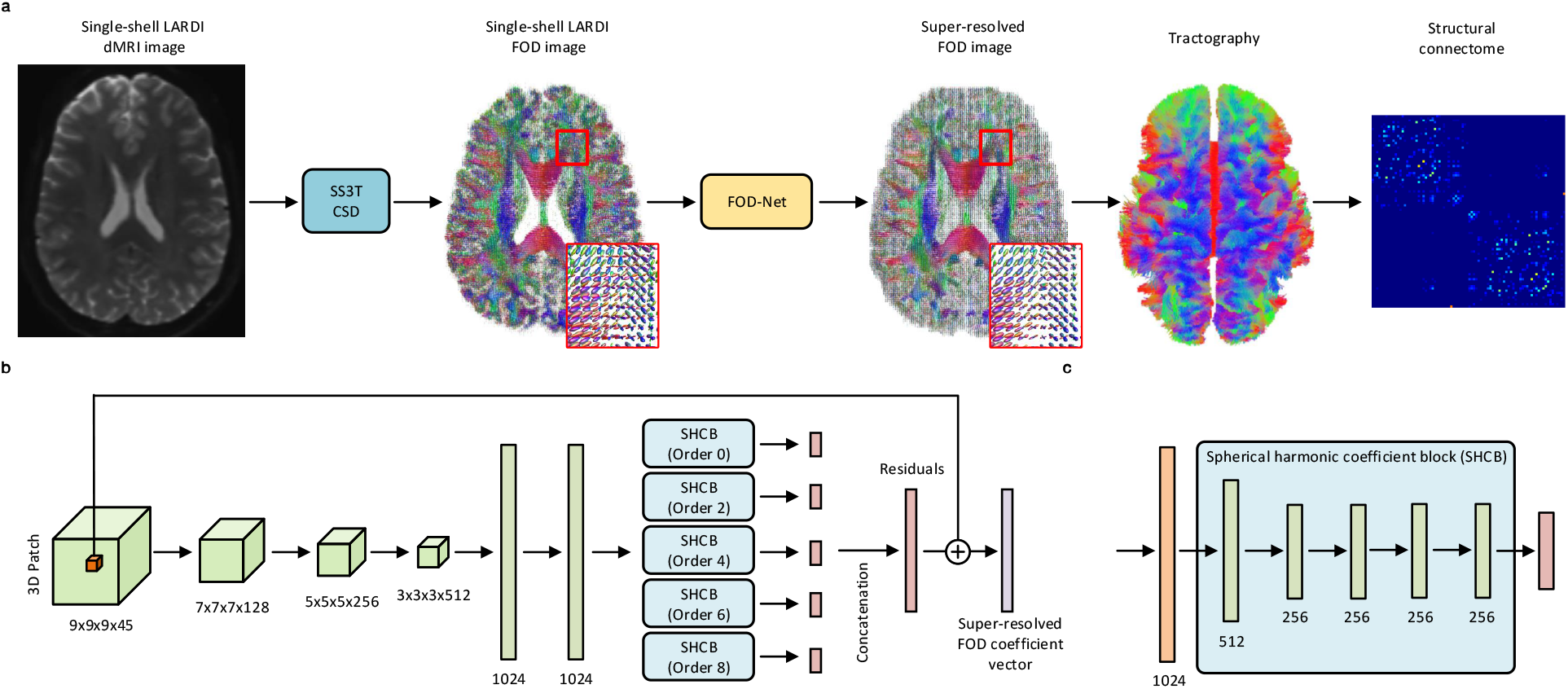
Schematic view of our fiber orientation distribution (FOD) network (FOD-Net) in a clinical connectome reconstruction pipeline and its network architecture. a, FOD-Net is flexible and can be embedded into a clinical connectome reconstruction pipeline to generate super-resolved FOD images, high-quality tractograms, and ultimately reliable structural connectomes. Super-resolved images are generated based on single-shell low angular resolution diffusion imaging (LARDI) derived FOD images, computed from single-shell 3-tissue (SS3T) constrained spherical deconvolution (CSD). b, The FOD-Net architecture consists of five convolutional layers, two fully connected layers and five spherical harmonic coefficient blocks (SHCB). It takes as input 4D SS3T CSD derived FOD patches and outputs the super-resolved version of the central voxel of the input patch. c, The architecture of spherical harmonic coefficient block, consisting of five two-dimensional convolutional layers.

FOD-Net consists of two stages: (1) the single-shell LARDI FOD patch feature representation stage, and (2) the superresolved central FOD voxel estimation stage. The feature representation stage is designed for extracting the feature vector which can represent the single-shell LARDI FOD patch 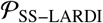 input implicitly. Subsequently, the output of the feature representation stage forms the input to the fiber orientation voxel estimation stage to obtain the coefficients of the spherical harmonic expansion of the super-resolved version 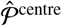 of the centre voxel of 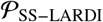. Both stages are trained jointly in an end-to-end manner.

The feature representation stage is composed of four 3D convolutional layers and two fully connected layers. All the 3D convolutional kernels of the 3D convolutional layers are of size 3, if not explicitly specified. Each 3D convolutional layer is followed by a batch normalisation layer and a gated linear unit (GLU) activation layer. The batch normalisation layers are used to normalise the output of the 3D convolutional layers by recentering and re-scaling, thus improving the stability of the training process. The first 3D convolutional layer, which has 128 filters, takes as input 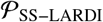. Then it is followed by four 3D convolutional layers with 256, 512, 1024, and 2048 filters, respectively (see Fig. 1b). All these four 3D convolutional layers do not use the zero-padding operation, and the size of feature maps outputted by each layer is therefore 7 × 7 × 7 × 128, 5 × 5 × 5 × 256, 3 × 3 × 3 × 512, and 1 × 1 × 1 × 1024, respectively. since the width, height, and length of the last feature map are all 1, this feature map is squeezed to the feature vector, which represents 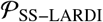. This feature vector is ingested by different spherical harmonic coefficient estimation blocks (SHCEBS) to estimate the residual between the coefficients of each order, respectively (see Fig. 1b). A SHCEB takes as input the feature vector, which is then processed progressively via a series of fully connected layers to obtain the coefficients of each specific spherical harmonic order. Since FODs are symmetric, the odd orders of the spherical harmonic coefficients are all 0, and only even spherical harmonic coefficients (five orders in our case) need to be estimated. Therefore, five sHCEBs are attached to the feature representation stage to compute the coefficients of each order of the central FOD voxel respectively. The size of each SHCEB output is 1, 5, 9, 13, and 17, respectively (Fig. 1c). The insights behind the design of a SHCEB come from two aspects. First, coefficients at each order have different distribution from those at other orders: while they have been all *z*-score normalised, the value range of coefficients is still different for each order. Thus, estimating coefficients using different blocks can significantly mitigate this issue. Second, spherical harmonics are orthogonal to each other, making it very difficult to predict all coefficients using the same vector. This motivates us to design a way to predict them separately. The final output of FOD-Net is the super-resolved FOD 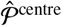 with 45 coefficients, which correspond to the super-resolution version of the centre voxel 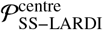 of 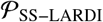. Since we are solving a regression task, the loss function to train FOD-Net is defined as:

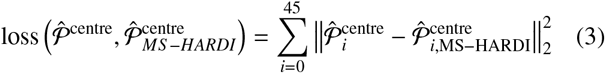

where 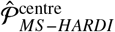 is the ‘ground truth’ multi-shell HARDI FOD supposed to be obtained from performing MSMT CSD on much more sophisticated dMRI acquisitions, *i* is the index of the coefficient of the spherical harmonic basis.

Existing deep learning models Lin et al. (2019); Tian et al. (2020) for dMRI angular super-resolution tasks are based on fixed-size dMRI images, which is not in principle suitable for our fiber orientation map angular super-resolution task because the size of dMRI data can vary considerably between acquisition protocols. To address this, the proposed FOD-Net takes (sequentially) as input an FOD patch cropped from a singleshell LARDI FOD image 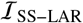 and outputs the angular super-resolved version of the central voxel of this patch. This formulation has two benefits: (1) FOD images with various sizes are able to be enhanced via the same trained FOD-Net model; (2) the use of a low angular resolution FOD patch for the central voxel angular super-resolution incorporates useful neighbor information given that the directions of FODs in a region are likely to be relatively consistent and, as such, this kind of information can compensate for lost angular information in LARDI. Although the larger patch size we choose, the more information we can include, we selected a 4D FOD input patch of size 9 × 9 × 9 × 45, as tradeoff between increase computational complexity and gain brought by larger patch sizes. 45 is the number of coefficients of spherical harmonic basis to represent an FOD. Furthermore, larger patch sizes lead to include more ‘noise’ (for example, unrelated information brought by grey matter voxels) from distant voxels, which may adversely affect the model performance.

We have implemented our system using Pytorch Paszke et al. (2019). The full network is trained in an end-to-end fashion from scratch, where all weights are initialized using the Xavier initialisation Glorot and Bengio (2010). Stochastic gradient descent (SGD) is employed as our optimizer because it generally acts well for regression tasks and has a favorable convergence speed. The parameters used for SGD are set to the defaults, with momentum 0.9. We use a learning rate of 0.01 with the learning rate decay 10^−5^ every epoch. The batch size is set to 64. The model has finished when training reaches around 20,000 epochs. One batch takes approximately 2s on a NVIDIA Tesla V100 GPU, and about two weeks for the model to converge.

In the inference stage of FOD-Net, single-shell LARDI FOD images were generated using the current state-of-the-art spherical deconvolution method for SS3T CSD, and then these FOD images were zero-mean unit-variance normalised using the statistics computed from the training dataset. A given normalised test image is sequentially cropped into 9 × 9 × 9 threedimensional (3D) patches, that are successively ingested by FOD-Net to obtain the super-resolved version of the central voxel of each input patch. Once the super-resolved FOD voxel for each matched single-shell LARDI FOD patch has been obtained, they are combined together back to generate the superresolved FOD data 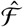.

### 2.3. Datasets

#### 2.3.1. The HCP dataset

The human connectome project (HCP) dataset is selected as the training dataset for our deep learning model due to the fact that it is one of the largest dMRI datasets in the neuroimaging community acquired using sophisticated HARDI protocols Sotiropoulos et al. (2013), which can be used to emulate the clinically accessible dMRI data via adequate subsampling operations, while still retaining the full high-quality dataset as ‘ground-truth’. Then the FOD data obtained from the subsampled dMRI (referred to here as single-shell LARDI) and the corresponding FOD data from multi-shell HARDI respectively constitute the low angular resolution and high angular resolution pairs for the training of our model.

The minimally pre-processed diffusion MRI data Sotiropoulos et al. (2013) in the HCP was acquired at a resolution of 1.25 mm isotropic, with 3 shells of *b* = 1000,2000,3000 s/mm^2^ (90 directions / shell) Van Essen et al. (2013). In addition, 18 *b*0 volumes were also acquired, leading to an overall dMRI data of 288 volumes for each subject. Since singleshell 32-direction dMRI data with low *b*-value is widely used in practical clinical settings, we generated clinically accessible dMRI data (i.e. single-shell low-angular-resolution) from the original multi-shell high-angular-resolution using the following subsampling strategies: we subsample 32 directions from the *b* = 1000 s/mm^2^ shell by selecting the first 32 directions sequentially from the original HARDI dataset, as the HCP acquisition pipeline samples directions in an incremental way such that an interruption at any point would result in an approximately optimal design Caruyer et al. (2011). Finally, the *b*0 volume is added into the extracted 32 directions to obtain the complete single-shell LARDI data (consisting of a total of 33 volumes for each subject).

Once clinical quality MR images and the corresponding multi-shell HARDI images are compiled, the next step is to generate their fiber orientation maps separately, using MRtrix Tournier et al. (2019). First, the Dhollander algorithm Dhollander et al. (2016) is used to compute the white matter response functions for all subjects’ data. Then, single-shell three-tissue (SS3T) Khan et al. (2020) constrained spherical deconvolution (CSD) and multi-shell multi-tissue (MSMT) Jeurissen et al. (2014) CSD are performed on the clinical quality MR images and multi-shell HARDI images to generate low angular resolution and high angular resolution FOD maps, respectively, by taking as input the computed white matter response functions. Subsequently, the multi-tissue informed intensity normalisation Raffelt et al. (2017) is performed on all the fiber orientation maps in the log-domain, which further corrects simultaneously for both global intensity difference as well as bias field. This step is necessary because it enables quantitative FOD comparison across subjects Raffelt et al. (2017). An intensity-normalised FOD dataset is four-dimensional, of which the first three dimensions are the spatial brain dimensions, and the last dimension is the coefficient volumes of the spherical harmonics expansion Tournier et al. (2004). In our experiments, the maximum harmonics order (*l_max_*) is set to 8, corresponding to 45 terms in the FOD spherical harmonic expansion, since it provides sufficient accuracy while remains low computational complexity Tournier et al. (2013). For each intensity-normalised FOD map, the mean and standard deviation of each coefficient volume are computed. Then the computed means and standard deviations are averaged across all the subjects to get the final means and standard deviations. Subsequently, the final means and standard deviations are used to perform *z*-score standardization on the intensity-normalised FOD maps to generate the standardized FOD maps. These standardized LARDI and HARDI FOD maps are ready for use in the training and inference stage of the FOD-Net.

110 subjects from the HCP dataset are randomly selected to build our dataset for conducting the experiments. Our dataset is split into 50 training, 50 evaluation, and 10 validation. The validation subjects are used for hyperparameter searching during the training process. Once the best hyperparameters of the network have been selected, all 50 subjects will be used to train the deep learning model.

#### 2.3.2. Local Clinical dataset

We conducted experiments with dataset acquired at our local clinical 3.0T scanner to further demonstrate the following two related key points. First, the good generalisation ability of FOD-Net, enabling assessing our model trained by HCP to be tested on data from a completely different clinical scanner/protocol. Second, to test the trained model on data from a more traditional clinical scanner and protocol (note that the Connectome HCP scanner is not a conventional clinical scanner, and thus the protocol used for the HCP dataset cannot be reproduced in standard clinical scanners). The local validation dataset was acquired using a 3.0T GE MR750 (General Electric, Milwaukee, WI, USA) with Nova 32-channel head coil. The multi-shell diffusion imaging parameters were: 70 axial slices, FOV 256 × 256 mm, TR/TE = 4000/82.3 ms, 2 mm slice thickness, acquisition matrix 128 × 128 with a 2 mm in plane resolution, and an acceleration factor of 2. The dMRI protocol included 136 diffusion-weighted images (32/32/72 directions of *b* = 1000/2000/3000 s/mm^2^) and eight *b* = 0 images. The acquisition time for the diffusion dataset was approximate 11 minutes. Additionally, a reversed phase encoding *b*0 “blip down” was obtained for susceptibility induced EPI distortion correction. An isotropic 1 mm T1-weighted image were also acquired using fast spoiled gradient echo (FSPGR) with magnetization-prepared inversion recovery (IR) pulse (TE/TI/TR = 2.62/450/7.06 ms, Flip Angle = 12).

To assess the generalizability, the aforementioned acquisition protocol was carried out on four healthy subjects (2 males, 2 females; 30 ± 5 years), in two occasions (for test-retest analysis) over a 30 mins session. The first scan of each subject was used to perform generalizability test. To assess intra-subject reliability, test-retest experiments were conducted on two difference timepoints of each subject.

### 2.4. Experiments

The comparison between FOD-Net and the state-of-the-art single-shell 3-tissue constrained spherical deconvolution (SS3T CSD) has been thoroughly made in three perspectives, including FOD accuracy, tractography qualitative performance, and connectome reconstruction accuracy, generalizability and test-retest reliability.

#### 2.4.1. FOD accuracy measurement

Qualitative and quantitative experiments are conducted to measure the FOD estimate accuracy of FOD-Net and SS3T CSD, respectively. Qualitative results are demonstrated in crossing-fiber regions of a randomly selected subject in HCP dataset (see Section 3.1). Furthermore, We quantitatively assessed the performance of our method on voxels containing pure white matter and in voxels containing a mix of white matter and grey matter (see Fig. 3), to assess whether the presence of GM contribution to the voxel dMRI signal impacted the performance of FOD-Net. We classified voxels in which the volume fraction of white matter is greater than 70% as pure white matter voxels. For voxels in the boundary between white matter and subcortical or cortical grey matter, we define them as partial volumed white matter voxels, as long as each of them contained at least 30% of white and subcortical/cortical grey matter tissues. The quantitative metric, i.e., angular correlation coefficient Anderson (2005), used to conduct both pure and partial white matter experiments can be calculated using:

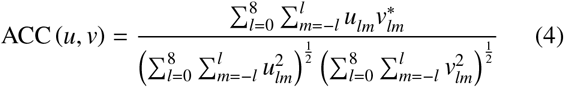

where *l, m* represents a degree and order of a spherical harmonic function, respectively, and *u, v* are defined as the ‘ground-truth’ FOD (MSMT CSD) and the estimated FOD data (SS3T CSD or FOD-Net), respectively.

The pure / partial white matter (a mix of white matter and grey matter) tissues are obtained using white and grey matter tissue segmentation provided in MRTrix Tournier et al. (2019); Smith et al. (2012), which generates a so-called ‘five-tissue-type’ mask for each subject, including cortical grey matter, sub-cortical grey matter, white matter, cerebrospinal fluid, and pathological tissue – Note: for our healthy subject data, the fifth tissue type was empty. These generated five-tissue-type masks are also used to perform the anatomically constrained tractography as illustrated in Section 2.4.2.

#### 2.4.2. Anatomical tractography reconstruction accuracy

To assess the tractograms generated by the super-resolved FOD images qualitatively (Fig. 4) and quantitatively (Fig. 5) in representative anatomic regions that might be of interest for clinical applications, we first segmented FOD data into different anatomical parts using TractSeg software Wasserthal et al. (2018), which is a deep-learning method for automatic white matter bundle segmentation based on FOD data. The bundles of interest included corpus callosum (CC), middle cerebellar peduncle (MCP), corticospinal tract (CST) and superior longitudinal fascicle (SLF). Then a bundle-specific tractography algorithm Wasserthal et al. (2019) was used to reconstruct anatomical tractograms from the segmented anatomic regions. Subsequently, three anatomical ROIs are defined to demonstrate the performance of FOD-Net on regions containing one-fiber-tract, two-fiber-tract, and three-fiber-tract, respectively. Specifically, ROI 1 is defined as a corpus callosum region, where is mainly comprised of large and coherent fiber tracts; we therefore use voxels only containing one fixel Raffelt et al. (2012) (a specific fiber bundle within a specific voxel) to evaluate our method in coherent fiber tract (non-crossing-fiber) regions. ROI 2 is defined as the regions where the MCP and CST bundles intersect, both of which can be regarded as fiber tracts possessing coherent orientations; the performance of FOD-Net on two-fiber-tracts-crossing regions can be evaluated using voxels in these intersected regions containing only two fixels. Regarding evaluations of regions with more complex fiber configurations (regions containing voxel with three distinct fixels), the intersected regions of SLF, CST, and CC are selected to conduct comparison experiments. Three metrics, including peak error, apparent fiber density error, and mean angular error, are selected to measure the fiber tract reconstruction accuracy in terms of these three ROIs. Peak and apparent fiber density (AFD) error can be computed using:

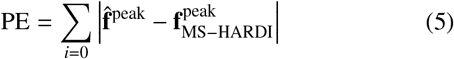

and

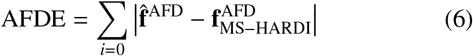

respectively. **f** and 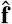 denotes a fixel estimate (generated from SS3T CSD or FOD-Net) and its corresponding ‘ground truth’ respectively. *i* is the index of a fixel within a voxel. Mean angular error (MAE), computed from the angular discrepancy between estimated and ‘ground-truth’ fixels, is defined as:

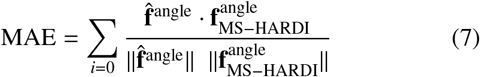

where **f**^angle^ is the direction vector which is of size 3, · is the dot product, and ||·|| is the *l*_2_ norm.

#### 2.4.3. Connectome reconstruction accuracy

To thoroughly evaluate the performance of super-resolved FOD images generated from FOD-Net, we compared the connectome reconstruction results between FOD-Net and SS3T CSD. The process of human connectome construction for a subject used in our connectomic analysis comprises five main steps: (1) whole-brain tractography; (2) computing a weighting for each streamline to make connectivity results quantitative Smith et al. (2015b); (3) grey matter atlas selection; and (4) connectome construction by computing the degree of structural connectivity between each pair of nodes. In the first step, anatomical-constrained whole-brain tractography is performed on all subjects using the HCP tractography pipeline presented in MRTrix Tournier et al. (2019). Specifically, by taking as input a given FOD image, the second-order integration over fiber orientation distributions (iFOD2) is used to compute the tractogram under the framework of anatomically-constrained tractography24 by providing the corresponding five-tissue-type mask. Relevant parameters were as follows: step size 0.625mm (0.5× voxel size), maximum curvature per step 45°, length 5 – 250 mm, and FOD amplitude threshold 0.06. Seeding operations are randomly performed from the interface between gray matter and white matter and 10 million streamlines are generated. For each tractogram, the SIFT2 Smith et al. (2015b) algorithm is used to determine a weight assigned to each streamline to make the tractogram quantitative and with more biologically meaningful properties. Subsequently, each subject’s anatomical image was segmented into biologically meaningful brain nodes using the Desikan-Killiany atlas in step 3. The fourth step isto construct the connectome matrix for each FOD-based tractogram: for each connectome reconstruction, streamlines were mapped to the relevant brain nodes using MRtrix, which takes as input the tractogram, the streamlines weights, and the atlas parcellation, and outputs the connectome matrix from the sum of the SIFT2 weights of all the streamlines (as measure of ‘connectivity’) connecting each pair of nodes in the parcellation.

The connectome disparity matrix, to evaluate the difference between a single-shell LARDI based connectome (estimates) and the corresponding multi-shell HARDI based connectome (‘ground-truth’), was computed as follows:

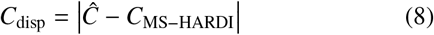

where 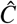 is a connectome matrix estimate computed by FOD-Net or SS3T CSD and *C*_MS–HARDI_ is the ‘ground truth’ connectome matrix.

#### 2.4.4. Generalizability test

To show how FOD-Net can adapt properly to new, previously unseen clinical acquisition protocols, the generalizability of FOD-Net was tested using the local clinical dataset (see above) acquired at our imaging centre. Since our model is trained using HCP data, in which each dMRI image is acquired at 1.25 mm isotropic resolution, we resized each image in our clinical dataset from 2 mm to 1.25 mm to match the configuration of our model. The resize operation is performed using cubic downsampling in MRtrix. Once all dMRI images had been resized, the single-shell LARDI dMRI image dataset was constructed by extracting all gradient directions in the first shell (*b* = 1000 s/mm^2^) and the *b* = 0 s/mm^2^ image. Next, the state-of-the-art SS3T CSD method was applied to all single-shell LARDI dMRI images to generate the single-shell LARDI FOD images. Then these LARDI FOD data were enhanced by FOD-Net to obtain the super-resolved FOD data. We also applied MSMT-CSD to the original multi-shell HARDI dMRI local clinical data to be used as ‘ground-truth’. Finally, tractography and connectome were reconstructed using the same processing pipeline as that described in the training stage.

We conducted two experiments to demonstrate the generalizability of our method. First, we computed the disparity matrices between the ‘ground-truth’ and our estimates using the absolute difference operator. Second, permutation testing was used to test whether the edges in the connectomes reconstructed using the super-resolved FOD images significantly differed from the ‘ground-truth’. Specifically, the null hypothesis of the equality in the distribution of the ‘ground-truth’ (MSMT CSD of HARDI) and the LARDI estimates (FOD-Net or SS3T CSD) was assessed using permutation testing, where each edge in the zero-diagonal and symmetric connectome matrices are processed independently. Then false discovery rate correction was applied to the results of the permutation test to ensure that corrected p-values are obtained. The null hypothesis was rejected if the tail of the distribution of an edge was longer in the estimate compared with the MSMT CSD based connectome (p value < 0.05).

#### 2.4.5. Test-retest reliability

Measuring test-retest reliability is of great importance because good test-retest reliability ensures the measurements obtained in one sitting are both consistent and stable, and that any improvement in accuracy introduced by FOD-Net does not come at the expense of a severe impact to its precision. In our study, the test-retest reliability measurements are conducted on all three methods (SS3T CSD, FOD-Net, and MSMT CSD) using the local clinical dMRI data acquired consecutively at two different timepoints. Specifically, for each timepoint, the single-shell LARDI dataset and multi-shell HARDI dataset are constructed using the same way as that in generalizability test. Then SS3T CSD and MSMT CSD are applied to the two single-shell LARDI and two multi-shell HARDI datasets, respectively. For the FOD images generated by SS3T CSD, we applied FOD-Net to generate the corresponding super-resolved FOD images. The connectome reconstruction pipeline which uses the SIFT2 weighted number of streamlines in MRTrix was performed for each FOD dataset generated by SS3T CSD, FOD-Net, and MSMT CSD. In test-retest experiments, the reliability of a method is measured by computing the average of the weighted mean coefficient of variation (wmCoV) Smith et al. (2015a) across the four subjects. wmCov is a metric to provide a summary statistic reproducibility of the entire connectome, which can be calculated by:

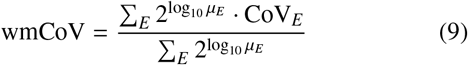

where CoV_*E*_ = *μ_E_*/*θ_E_* is the coefficient of variation (CoV) of edge *E*; *μ_E_* and *θ_E_* are the mean and the unbiased standard deviation estimate of mean connectivity of edge *E* across the connectomes reconstructed from a same subject (intra-subject). wmCoV computes a representative mean CoV for a group of multiple scans on a same subject, where the contribution of each edge CoV to the mean is weighted using the mean connectivity of that edge.

## 3. Results

### 3.1. FOD accuracy

To assess the quality of FOD angular super-resolution, we compared the FOD-Net predictions with the ‘ground-truth’ (obtained by conducting multi-shell multi-tissue (MSMT) constrained spherical deconvolution (CSD) on the multi-shell HARDI data6) and with the single-shell LARDI FOD data computed from the current state-of-the-art method (i.e. single-shell three-tissue (SS3T) CSD Dhollander et al. (2016)). A randomly selected LARDI FOD image containing two anatomical regions of interest (ROI) is shown Fig. 2 a. Fig. 2 b-d and e-g show the two zoomed-in regions of interest, revealing 3D FODs associated with crossing fiber tract regions. FOD-Net was applied to single-shell LARDI FOD data of subjects (i.e., test data) previously unseen by the network during the training process, with the resulting super-resolved FOD images shown in Fig. 2c,f: the complex fiber architecture features are clearly resolved by FOD-Net, and the FODs show consistent agreement with the ‘ground-truth’ FOD data shown in Fig. 2d,g. Furthermore, quantitative assessment of FOD-Net using the angular correlation coefficient (ACC) metric (Fig. 3) approximated the ‘ground-truth’ when applied to both pure white matter voxels and to voxels of mixed gray-white matter type (See addition scores in Table 1). We can found that FOD-Net provides a robust method for FOD enhancement.

**Fig. 2.**
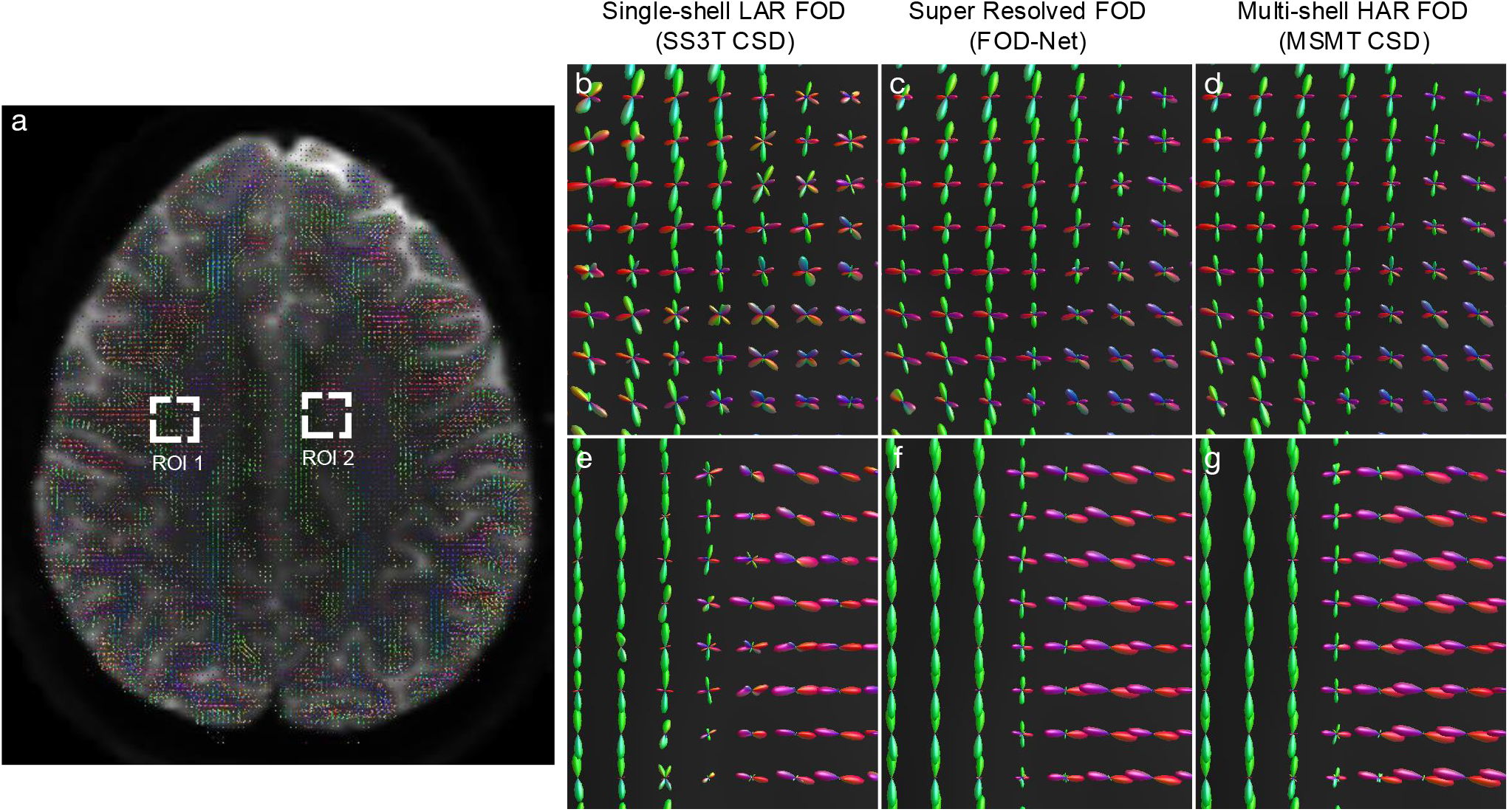
Super-resolved fiber orientation distribution (FOD) images using FOD-Net. a) Single-shell LARDI FOD image generated from single-shell LARDI data using single-shell 3-tissue (SS3T) constrained spherical deconvolution (CSD). Two regions of interest (ROIs) are magnified and shown in b-d (ROI 1) and e-g (ROI 2), respectively. (b,e), (c,f), and (d,g) are single-shell LARDI, single-shell LARDI super-resolved, and multi-shell HARDI ‘ground-truth’ FOD images generated using SS3T CSD, FOD-Net, and multi-shell multi-tissue (MSMT) CSD, respectively. Similar findings were obtained in the Human Connectome Project test data and our local 3.0T clinical dataset.

**Fig. 3.**
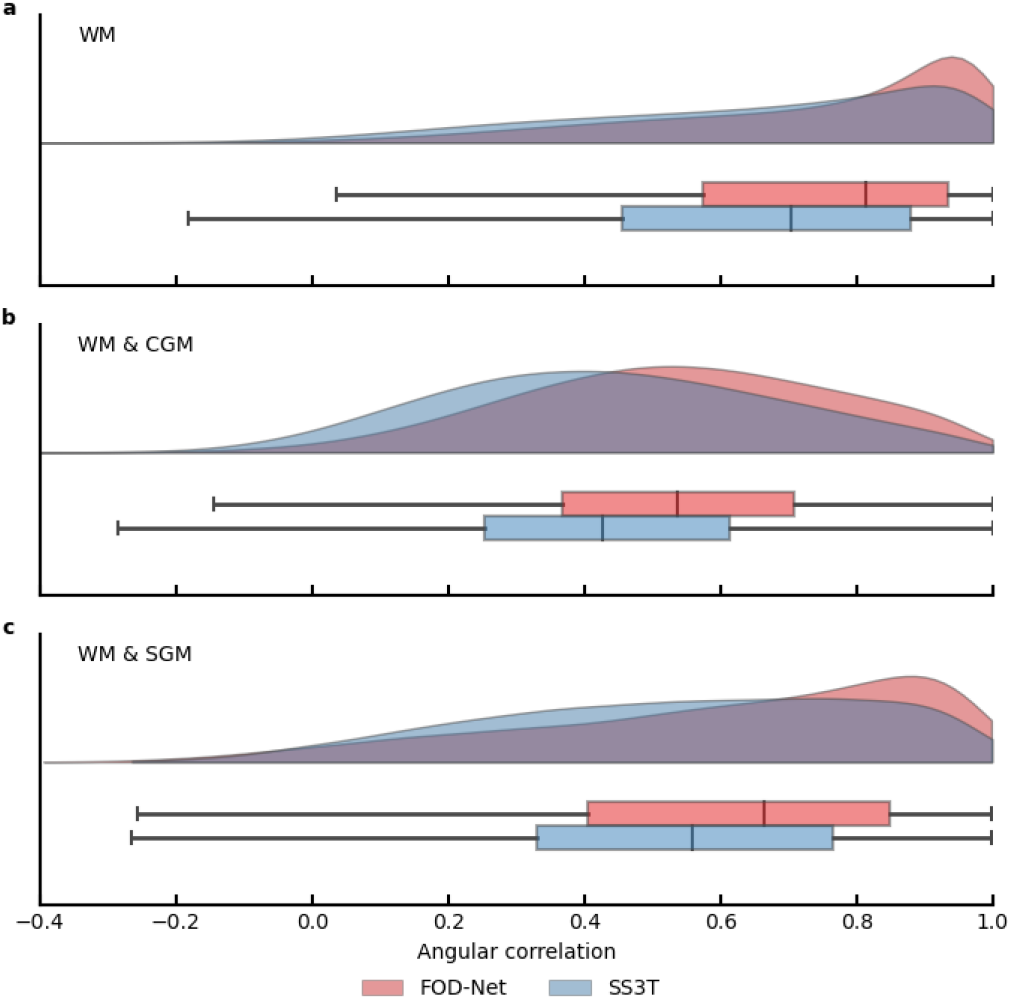
FOD-Net provides more accurate FOD estimates compared to SS3T CSD in both pure white matter (WM) and partial volumed white matter voxels. a-c) inference results of 50 subjects from the Human Connectome Project test set, measured by the angular correlation coefficient (ACC) metric to the ‘ground-truth’. a) Pure WM tissue. b-c) Voxels of WM partial volumed with cortical grey matter (GM), i.e. in the interface between WM & cortical GM, and with subcortical GM (i.e. in the interface between WM & subcortical GM), respectively. Half-violin plots report the distribution of the ACC of the FOD estimates of each method. The statistical information, including the minimum, maximum, upper/lower half quartile, and mean values are shown in each boxplot. Compared with SS3T CSD, FOD-Net results are much closer to the ‘ground-truth’.

**Table 1.**
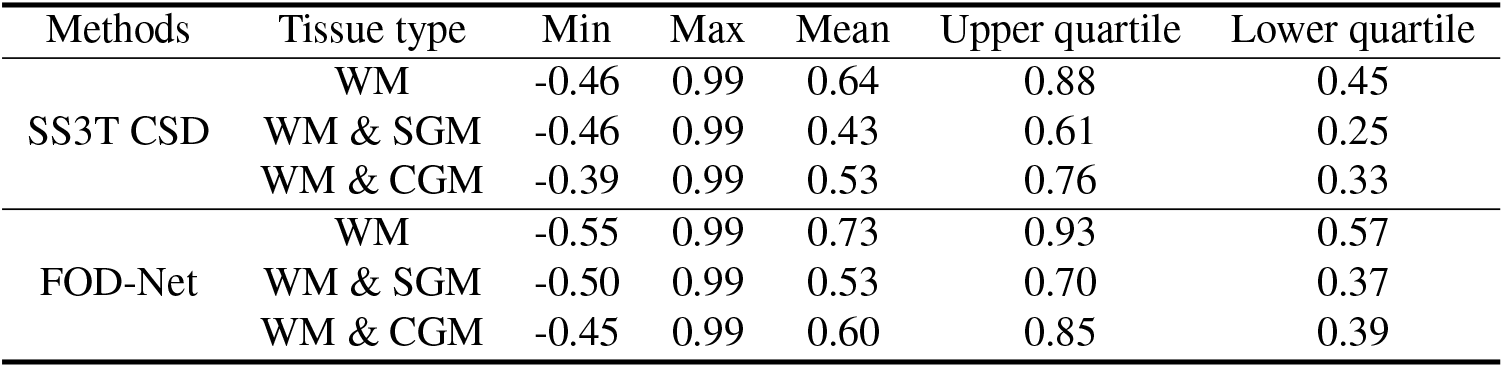
The evaluation results performed in both pure white matter and partial volumed white matter voxels using angular correlation coefficients (ACC) metric. Partial white matter regions are voxels of WM partial volumed with cortical grey matter (i.e. in the interface between WM & cortical GM) and with subcortical grey matter (i.e. in the interface between WM & subcortical GM) respectively.

### 3.2. Tractography in anatomical regions of interest

FOD-Net removes spurious fibers and recovers missing fiber tracts in anatomical regions of interest. To further demonstrate the resolution improvement and the flexibility of FOD-Net, we replaced SS3T CSD with FOD-Net in the tractography pipeline of MRtrix, a widely used dMRI software tool Tournier et al. (2019). Whole-brain tractograms are generated based on superresolved FOD data; then, three anatomical ROIs (See Section 2.4.2 for the definition of the three anatomical ROIs) containing representative fiber tract configurations are selected for qualitative comparison with SS3T CSD. Fig. 4 a-c illustrate corpus callosum fibers generated using SS3T CSD, FOD-Net, and ‘ground-truth’, respectively. In general, FOD-Net produced smoother, clearer and less noisy (see Fig. 4 a,b) fiber tracts than SS3T CSD, more closely corresponding to the configurations observed with the ‘ground-truth’. In crossing fiber regions (Fig. 4 d-f), tractograms generated based on our method produced denser fiber tracts (Fig. 4 e,h, respectively) that more closely aligned with the ‘ground-truth’ (Fig. 4 f,i) than SS3T CSD-derived FOD images (Fig. 4 d,g), in which spurious and missing (sparse) connections were frequently observed. We have also observed that the FOD-Net results are inclined to produce tractograms with smoother fiber tracts compared to the ground truth, especially for large single fiber bundles (Fig. 4 a-c). This result agrees with the neuroanatomical fact that a fiber tract and its neighbors are most likely to have a consistent direction and FOD-Net has gained this knowledge from the training data.

**Fig. 4.**
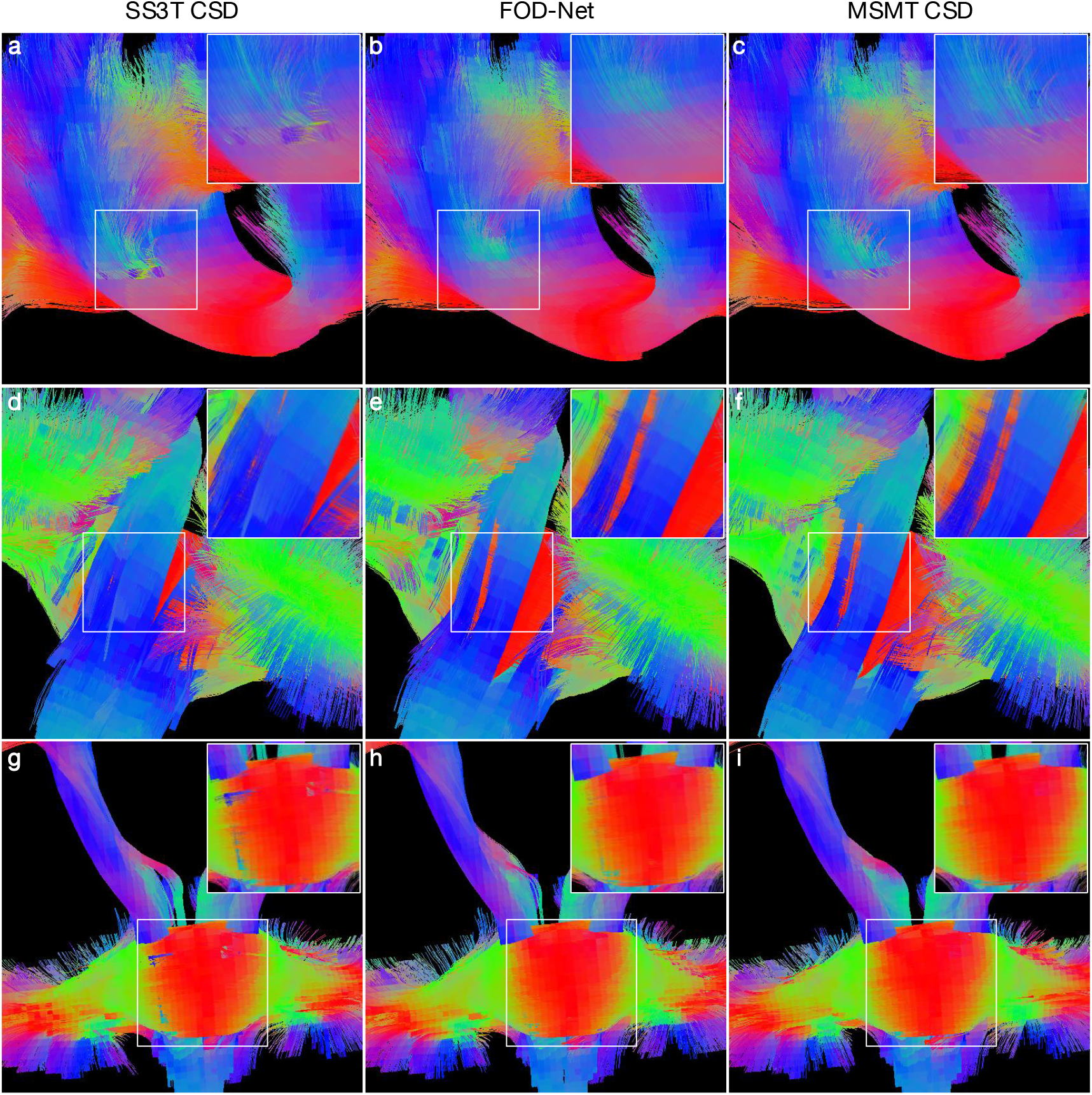
Tractography improvement by FOD-Net in three anatomical regions of interest. a-i) Examples of tractography generated based on SS3T CSD of LARDI (a,d,g), FOD-Net of LARDI (b,e,h), and MSMT CSD of HARDI (ground truth) (c,f,i), and magnified view of the boxed regions. In the boxed region of (a), FOD-Net resolves the irregular pattern of a large single fiber, in close agreement with the ground truth (c). (e,h) Reconstructed higher density fiber tracts in the fiber crossing regions, which match the results obtained from (f,i). However, (d,g) have sparser appearance and generate numerous spurious fiber tracts. Similar findings were observed with 50 subjects from the Human Connectome Project test set. The colour-coding indicates the local fiber orientation, with red: left-to-right, green: anterior-to-posterior, and blue: rostral to caudal.

In addition, quantitative results measured by the peak error Smith et al. (2015b), mean angular error (MAE), and apparent fiber density Raffelt et al. (2012) (AFD) error (Fig. 5 a-c) were used to assess quantitatively the quality of FOD estimation. For the tissues containing large, coherent fiber tracts (ROI 1, corpus callosum), our method achieved a peak error of 0.076±0.005 (mean ± SD) and an apparent fiber density error of 0.075 ± 0.006, while SS3T CSD attained 0.23 ± 0.04 (pj0.0001, p indicates two-tailed p value obtained from t-test to test if SS3T CSD results is statistically significant when compared to FOD-Net values) and 0.22 ± 0.04 (*p* < 0.0001), for these values, respectively. Furthermore, FOD-Net achieved similar results (peak error, 0.073 ± 0.005; apparent fiber density error, 0.062 ±0.004) on tissues containing two intersecting fiber tracts (ROI 2, intersected regions between middle peduncle and corticospinal tract); in this region, SS3T CSD achieved 0.12 ± 0.02 (*p* < 0.0001) and 0.11 ± 0.02 (*p* < 0.0001) respectively, which are lower by a large margin (around 60%). Finally, we report the FOD-Net results (peak error, 0.063 ± 0.008; apparent fiber density error, 0.051 ± 0.004) measured in three-fiber-tract crossing regions (ROI 3, intersected regions between the superior longitudinal fascicle, corticospinal tract, and corpus callosum), where SS3T CSD attained 0.097 ± 0.007 (*p* < 0.0001) and 0.081 ± 0.006 (*p* < 0.0001) respectively. Notably, FOD-Net outperformed SS3T at all three anatomical ROIs, which contained a range of representative fiber tract configurations. The cumulative distribution of mean angular error, computed from the angular discrepancy between estimated and ‘ground-truth’ fiber bundles, is shown for all three ROIs in Fig. 5 c. We observed that the percent of ROI-1 voxels with a MAE less than 10° was approximately 95% for FOD-Net and only 84% for SS3T CSD. By observing ROI-2 voxels with a MAE less than 10°, FOD-Net outperformed SS3T CSD by a large margin, i.e., the percent of voxels meeting the criteria increased from 35% to 65%. For ROI-3 voxels of which MAE is less than 10°, FOD-Net achieved significant improvement (10% to 50%) over SS3T CSD. Interestingly we found that FOD-Net yielded larger improvements in crossing-fiber regions (ROIs 2 and 3) than coherent single fiber regions (ROI 1), which indicates that ambiguities generated by a high number of crossing-fiber voxels in ROIs 2 and 3 were successfully resolved by FOD-Net. We also found that the FOD-Net outperformed SS3T CSD in terms of MAE in pure and partial white matter tissues (See Fig. 6).

**Fig. 5.**
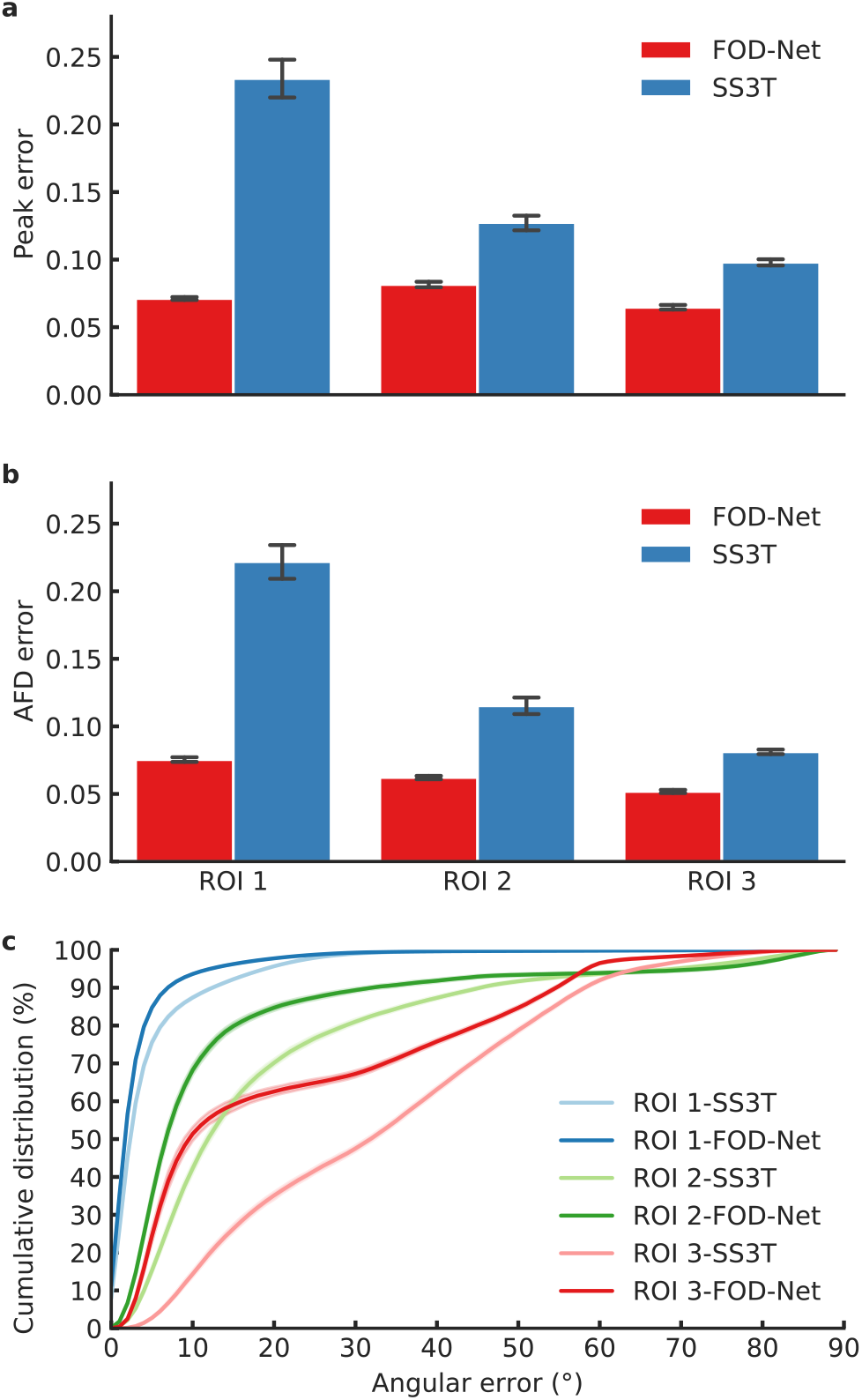
Average of voxel-wise quantitative results across 50 subjects from the test HCP dataset with respect to three anatomical regions of interest (ROI1: single fiber region; ROI2: 2-fiber crossing region; ROI3: 3-fiber crossing region). a) Peak error computed by the absolute difference between the amplitude of a FOD estimate at the maximum peak and its corresponding ‘ground-truth’. b) Apparent fiber density (AFD) error computed by the absolute difference between the integral of a given FOD estimate and its corresponding ‘ground-truth’. c) The cumulative diagram represents the fraction of the number of fiber tracts measured by the angular error.

**Fig. 6.**
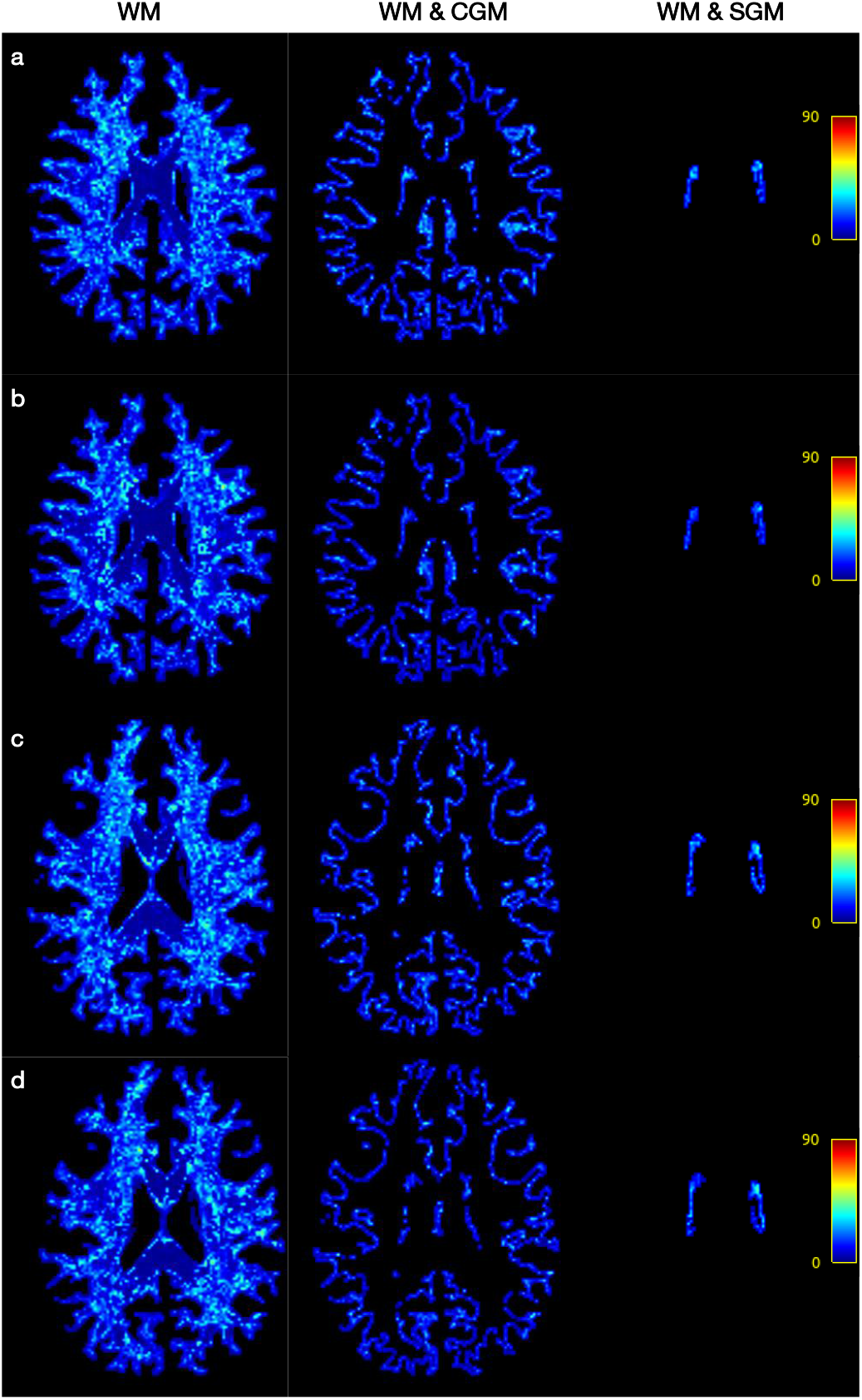
Mean angular error results measured on pure and partial white matter volumes (Please refer to Section 2.4.1 for pure / partial volume definition) of two subjects using SS3T CSD and FOD-Net respectively. a-b, mean angular error measured on a slice of a same subject by SS3T CSD (a) and FOD-Net (b) respectively, in terms of white matter (WM), the interface between white matter and cortical grey matter (WM & CGM), and the interface between white matter and subcortical grey matter (WM & SGM). SS3T CSD (c) and FOD-Net (d) results on a slice of another subject are shown.

### 3.3. Connectome reconstruction from FOD images

More reliable connectomes are generated by the enhanced fiber orientation images. One widely used application for dMRI tractography is the macro-scale brain structural connectome Seguin et al. (2019); Fornito et al. (2015, 2016). After performing whole-brain tractography from the super-resolved FOD data, the regions defined in the Desikan-Killiany cortical atlas Desikan et al. (2006) were employed as network nodes, and the tracks connecting these nodes were used to calculate the structural connectome.

We report the accuracy of connectome reconstruction, quantified using absolute error in terms of the connectivity number of streamlines (Fig. 7 a-c). Fig. 7 a (the first three columns) shows the average connectome matrices across healthy subjects generated by SS3T CSD from LARDI, MSMT CSD from HARDI, and FOD-Net from LARDI, respectively. Fig. 7 a (the last two columns) shows the absolute disparity map between the connectome matrix of MSMT CSD and the LARDI estimates (SS3T CSD or FOD-Net). We can observe that FOD-Net yielded the most accurate connectome reconstructions, outperforming SS3T CSD by a large margin. In particular, for the group-averaged connectomes, and using the MSMT CSD results as ‘ground-truth’, implausible connections (for example, lateral occipital and pericalcarine; posterior cingulate and rostral middle frontal; fusiform and inferior temporal; lingual and pars orbitalis; inferior temporal and pars orbitalis; etc.) and missing connections (for example, transverse temporal and insula; accumbens area and thalamus; rostral anterior cingulate and insula; etc.) from the conventional singleshell based method (SS3T CSD), are respectively rectified and ‘reconnected’ by FOD-Net (See Fig. 8 for brain node pairs with top 5 numbers of spurious and missing connections, respectively). Minor discrepancies between the FOD-Net and ‘ground-truth’ connectome are attributable to error conferred by low density connections (cuneus and fusiform; fusiform and lingual); and were also observed in the SS3T CSD-derived connectome. From Fig. 8, we can also observe that the performance of both SS3T CSD and FOD-Net is degraded in inferior temporal and middle temporal; and superior parietal and precuneus regions. However, FOD-Net still outperformed SS3T CSD in these regions significantly.

**Fig. 7.**
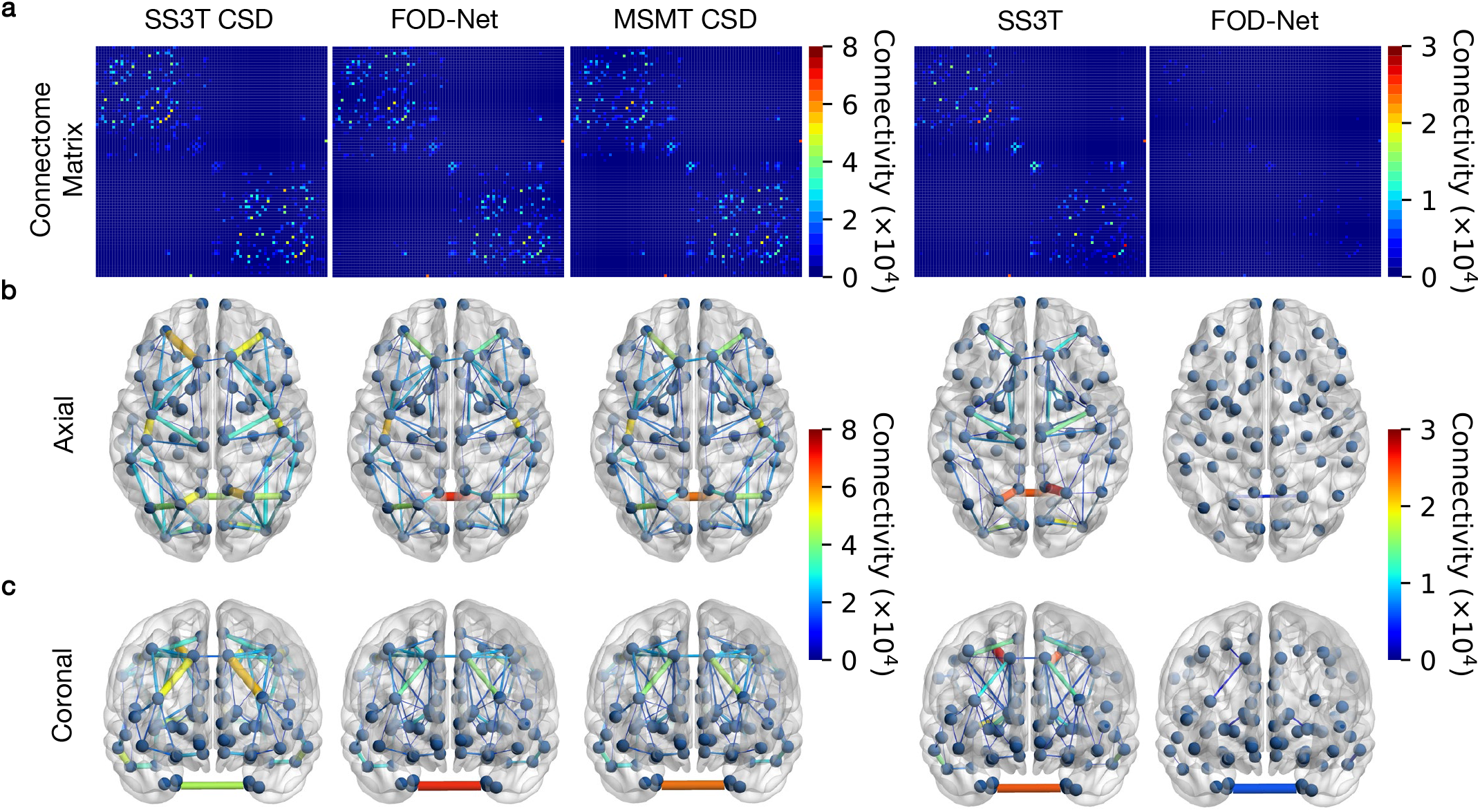
Average structural connectivity matrices across fifty healthy adults from the test HCP dataset. a, Averaged structural connectome matrices generated by SS3T CSD from LARDI, MSMT CSD from HARDI, and FOD-Net from LARDI, respectively. The last two columns show the disparity matrices between MSMT CSD (considered as ‘ground-truth’) and the LARDI estimates (SS3T CSD or FOD-Net), respectively. b-c, Results are also rendered in anatomical space using the axial and coronal view. To better visualize the main differences 240 between the ground truth and the estimates, only the connections of which the number of streamlines is larger than 8000 (10% of the upper bound of the colour bar of the connectome matrices in a) are shown in the first three columns. The connection disparity visualisation threshold is set to 3000 (10% of the upper bound of the colour bar of the disparity maps in a) for the last two columns.

**Fig. 8.**
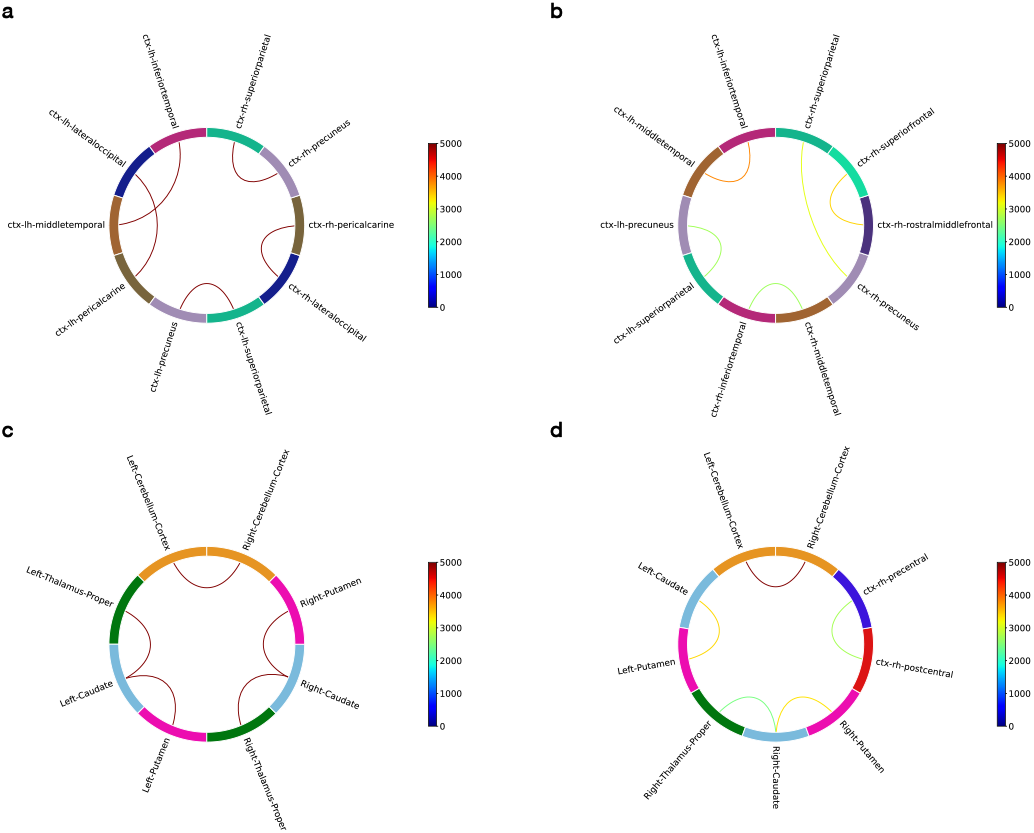
Implausible and missing connections (top 5 brain node pairs for each type of connections) generated by SS3T CSD and FOD-Net respectively. a-b, brain node pairs with top 5 numbers of implausible connections (false positive connections) of SS3T CSD and FOD-Net respectively. c-d, brain node pairs with the top 5 numbers of missing connections (false negative connections). The colours indicate the difference between an estimated connection (SS3T CSD and FOD-Net) with the ground truth one (MSMT CSD).

To further explore these results, we also examined the density of self-connections in the brain atlas, i.e. computed based on the streamlines connecting each node with itself; our results demonstrate that FOD-Net of LARDI data achieved more similar intra-node connectivity to MSMT CSD HARDI than SS3T CSD did (see Fig. 9). Fig. 9 a-c represents the density of self-connections of SS3T CSD, MSMT CSD, and FODNet, respectively, in the brain atlas. Fig. 9 d,e represents the fraction of the count of mispredicted connectivity strength (missing and spurious connections) in each region defined in the brain atlas. In cerebellum, we found that FOD-Net achieves more similar results with MSMT CSD compared to SS3T CSD. We also observed that SS3T CSD tends to produce a significant number of spurious fiber tracts in regions including rostral middle frontal, superior frontal, superior temporal, supra marginal, and superior parietal gyrus. The high proportion of these spurious connections in the corresponding brain regions could adversely affect the accuracy of the reconstructed connectome for clinical applications. In contrast, FOD-Net achieved superior performance in these clinically challenging brain regions. It is worth mentioning that many missing connections are found in SS3T CSD results of thalamus, caudate, and putamen, which contain complex fiber configurations. This is because SS3T CSD performed on single-shell LARDI has more difficulties at resolving fiber orientations reliably. Our method significantly reduces the number of missing connections by ‘reconnecting’ them through FOD super resolution. It should be noted that both FOD-Net and SS3T CSD make large errors in missing and spurious connections in the pallidum region. However, our method still achieves better results than SS3T CSD. The high proportion of SS3T CSD derived spurious intra-node connections in the corresponding brain regions could adversely affect the accuracy of the reconstructed connectome for clinical applications.

**Fig. 9.**
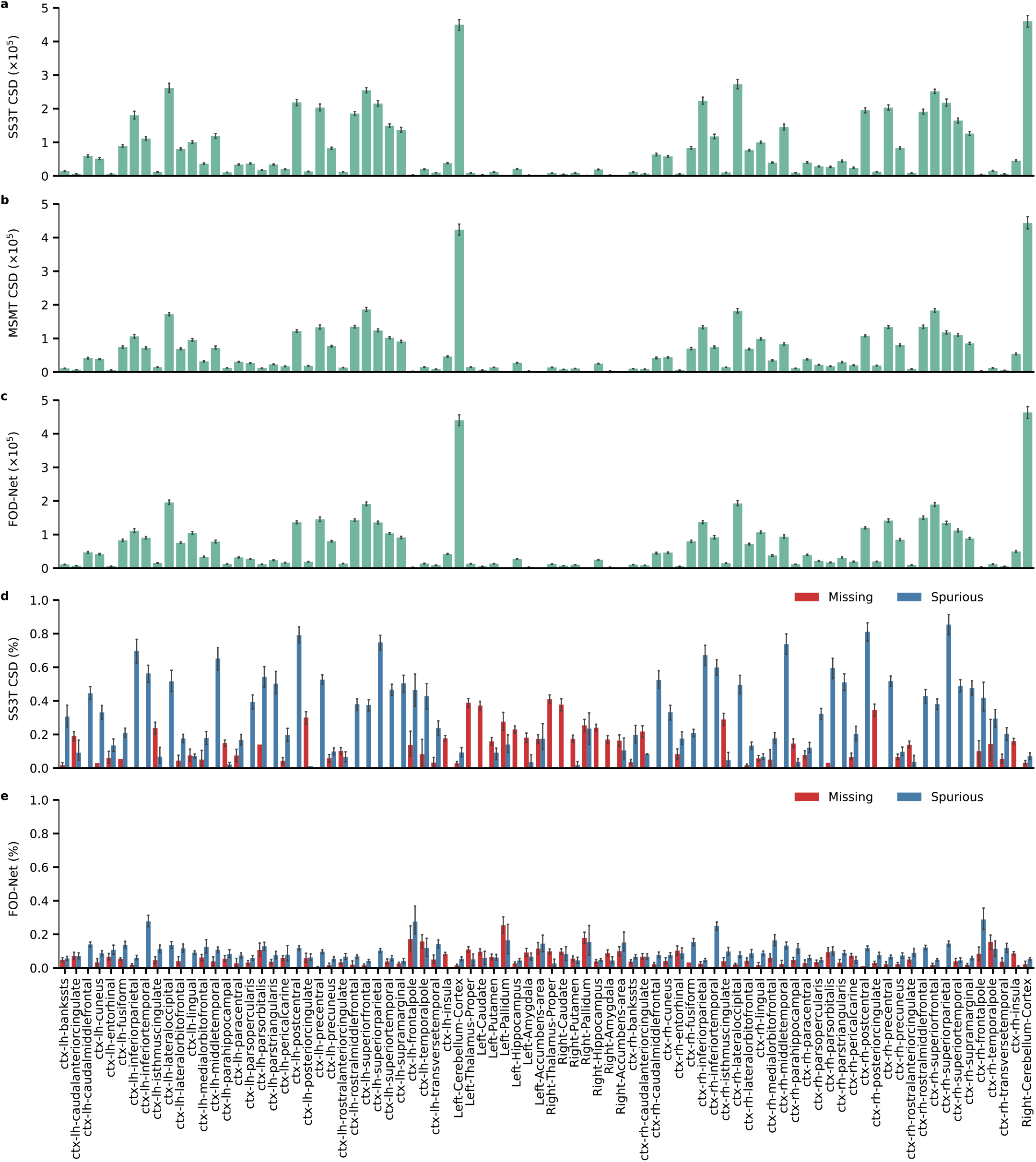
The density of self-connections in the brain nodes for the test HCP dataset. a-c, Summary of the density of self-connections in the brain nodes in terms of SS3T CSD of LARDI, MSMT CSD of HARDI, and FOD-Net of LARDI. d-e, Fraction of the number of missing and spurious streamlines when comparing MSMT CSD results with the LARDI estimates (SS3T CSD or FOD-Net).

Permutation testing was used to test whether the connectomes reconstructed using LARDI data significantly differed from the ‘ground-truth’ HARDI connectomes. Fig. 10 shows that over 74% connections generated by SS3T CSD (a) were statistically significant different from the ‘ground-truth’, suggesting that structural connectomes generated from single-shell LARDI FOD images are therefore likely to deviate from the ‘truth’ and fail to robustly model complex fiber tract configurations for existing clinical applications. In contrast, only 4% connections in the FOD-Net-based connectome show significant differences from the ‘ground-truth’, which demonstrates the superiority of super-resolved FOD data to reconstruct more reliable connectomes without the need for time-consuming multi-shell HARDI data.

**Fig. 10.**
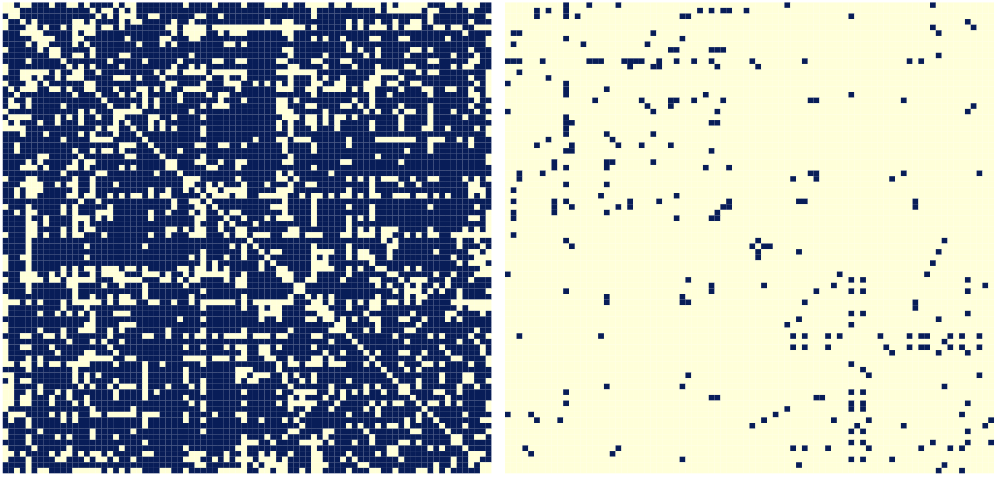
Statistical significance test between MSMT-CSD of HARDI based connectome matrices and LARDI based connectomes (blue indicates statistical significance found at *p* value < 0.05, false-discovery-rate-corrected) for the HCP test dataset. Significantly different edges for: a, SS3T CSD, b, FOD-Net. More than 74% connections in SS3T CSD (a) were statistically significant different compared with the ‘ground-truth’, while only 250 4% were for FOD-Net (b).

### 3.4. Generalizability and reproducibility

FOD-Net has good generalizability and reproducibility. We tested the generalizability of our trained deep learning model in improving FOD quality from clinical diffusion MR images acquired by unseen (not present in the training phase) protocols using a different clinical MRI scanner. From Fig. 11, improvement of connectome reconstruction obtained from superresolved FOD images of LARDI data is very consistent with the HCP results from Fig. 7 despite the clinical data having been acquired from protocols unseen by FOD-Net in the training stage. In contrast to FOD-Net, the connectome matrices based on SS3T CSD on LARDI data exhibit large amounts of false positive and false negative edges (compared with the ‘groundtruth’ MSMT-CSD HARDI results), hampering the reliability of clinical connectomics applications.

**Fig. 11.**
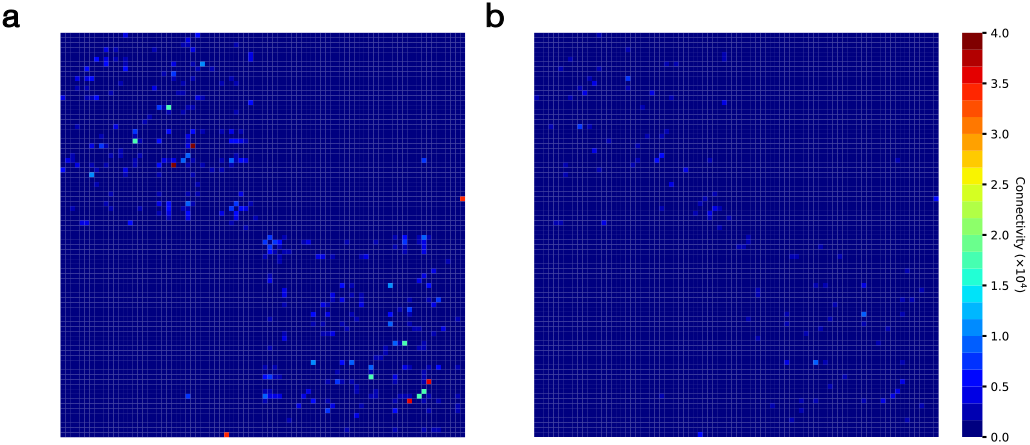
The connectomes based on super-resolved FOD images yield more robust and reliable connection reconstruction for unseen clinical protocols (not present in the model training phase). a) Disparity map computed by subtracting the SS3T CSD based connectome with the MSMT CSD based (‘ground-truth’) connectome. b) Corresponding disparity map for the FOD-Net based connectome. All connectome matrices are calculated by averaging across 4 clinical subjects.

We also examined the reproducibility of our method by performing test-retest experiments on data acquired by clinical protocols that differed from those used in the training data. We assessed the reproducibility of the structural connectomes generated by SS3T CSD, FOD-Net, and MSMT CSD, respectively, in the intra-subject scenario, with the connectome measured by SIFT2 connectivity. The intra-subject connectome difference for each FOD method therefore reflects the test-retest connectome reliability associated with a given FOD method. We computed weighed mean coefficient of variation (wmCoV) (a test-retest metric to provide a summary statistic of a group of connectomes) for each subject, then averaged these metrics across all subjects to yield final scores of 0.61 (SS3T CSD), 0.75 (FOD-Net), and 0.72 (MSMT CSD). One can observe that the reproducibility score of FOD-Net is very close to that of MSMT-CSD. While SS3T CSD achieved the best reproducibility (*p* = 0.02) in our clinical dataset by a small margin, FOD-Net connectome reproducibility was not statistically significant when compared to MSMT CSD (*p* = 0.79). Fig. 12 shows the absolute difference between test-retest connectomes for each FOD method.

**Fig. 12.**
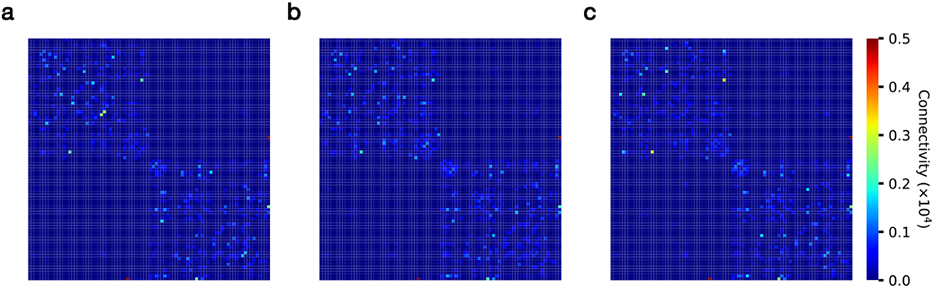
FOD-Net has good test-retest reproducibility for dMRI images acquired from same subjects at different timepoints. (measured by SIFT 2 connectivity). a-c, the disparity maps computed by SS3T CSD, MSMT CSD, and FOD-Net, respectively.

## 4. Discussion

The proposed deep-learning-based method, FOD-Net, facilitates the generation of angular super-resolved FOD images directly from conventional clinical-type single-shell LARDI data.

FOD-Net produces FODs of equivalent quality to those produced from more advanced multi-shell HARDI data, but requires only a fraction of the dMRI data (e.g. with as little as 11% of the dMRI data for the HCP examples used here). Furthermore, FOD-Net achieves these FOD improvements without requiring high *b*-value data, which is most often not available in clinical protocols, but has been shown to be required for optimal performance when using current FOD algorithms Tournier et al. (2013,2020). The relaxing of the requirement for high *b*-values also leads to an associated increase in SNR of the dMRI data, as higher *b*-values require longer TEs, in turn resulting in reduced SNR Jones et al. (1999). In addition to FOD angular superresolution, our method can be flexibly embedded into modern dMRI analysis pipelines for clinical research applications11,13. By exploiting super-resolved FOD images derived from clinical acquisitions, we demonstrate enhanced tractography applications including fiber tacking, attributable to fewer false positive and false negative connections in crossing fiber regions compared to the current state-of-the-art SS3T CSD method. Furthermore, we demonstrate that FOD-Net can produce structural connectomes from common clinical single-shell LARDI protocols of similar quality to those obtained with more sophisticated (but time-consuming) multi-shell HARDI protocols used in advanced research applications. This opens up the possibility for more reliable connectomics analysis from widely used clinical protocols (e.g. 32 directions, *b* = 1000 s/mm^2^), which are generally considered to be of insufficient quality for reliable analysis.

Algorithmically, the deep network takes as input a 3D patch cropped from a single-shell LARDI FOD image and outputs the super-resolved FOD voxel corresponding to the central voxel of the input 3D patch. By repeating this process on all 3D patches of a low angular resolution FOD image, the deep network generates all super-resolved FOD voxels, which can be combined to form up the corresponding super-resolved FOD image. Since our method is patch-based, any size of FOD images can be ingested by our model to output the corresponding super-solved images, giving FOD-Net great flexibility when dealing with data with different spatial resolutions. Another merit of this patch-based design is that information from neighboring voxels is incorporated to provide clear clues to compensate for the information lost in single-shell LARDI FOD voxels, given that anatomical white matter fiber tracts are likely to have consistent orientation with those in neighboring voxels. Importantly, our formulation is based on the use of FOD images, for which the size of the last volume axis can be fixed (i.e. by the choice of maximum harmonic order *l_m_ax*), which can circumvent the problem of constructing dMRI-based deep learning models from a range of acquisition protocols: while the numbers of gradient directions (corresponding to the size of the last volume axes in the dMRI dataset) can considerably vary between acquisition protocols, deep learning models typically require a fixed input size; the use of a fixed *l_m_ax* overcomes this limitation, thus facilitating the generalization of FOD-Net to varying acquisition protocols. Moreover, the superior performance achieved by FOD-Net is achieved by the regression of the residual between the single-shell LARDI FOD images and the corresponding super-resolved FOD images. This is because they share a high degree of mutual information, with an output probability distribution that is conditional upon the input single-shell LARDI FOD distribution. A necessary step before the use of FOD-Net is FOD normalization, which standardizes each dimension of a FOD to enhance the numeral stability of the network. While original magnitudes of the coefficients of the high order terms of the FOD spherical harmonics expansion are very close to zero, these sort of coefficients have significant impact on the quality of the FOD estimates, as they determine the overall angular resolution20. A proper normalization step can rescale the value range of these higher order terms, such that the network can be more influenced by them to enhance the performance and converge faster.

Despite FOD-Net providing more accurate FOD estimates than SS3T CSD in both pure white matter and partial volumed white matter voxels (see ACC metrics in Fig. 3), we found that the performance of FOD-Net in pure white matter voxels outperforms that in white matter voxels with partial volume with grey matter structures (see Fig. 3). This observation is in line with the fact that white matter voxels comprising partial volume with large amounts of grey matter, in which nerve cell bodies and dendrites do not have clear and coherent orientations, reduce the overall amplitude of the FOD Calamante et al. (2018). Nevertheless, connectomes reconstructed from super-resolved FOD images are still reliable and robust, as tracking is often terminated at the grey-white matter interface Smith et al. (2012).

With respect to tractography, Fig. 4 shows that by using super-resolved FOD images, the tractography results are improved compared with the current state-of-the-art SS3T CSD method, providing tractograms that reflect better those obtained with the ‘ground-truth’ multi-shell HARDI data. This likely reflects the relatively lower angular resolution of clinical dMRI protocols, which provide insufficient information for SS3T CSD to reliably infer the specific fiber architecture within the voxels. Furthermore, the use of low *b*-values in typical clinical protocols exacerbates this problem by reducing the angular contrast of the diffusion effect4,5, which limits the ability to characterize fine-grained fiber tract properties. Our deep learning strategy helps to bridge this gap, increasing the angular resolution and generating super-resolved FODs: unlike conventional SS3T CSD, FOD-Net learned the inherent relationship between single-shell LARDI and multi-shell HARDI FOD data, which means it can reveal the sophisticated intravoxel white matter architecture in fine detail. It is also worth noting that the output of FOD-Net is smoother, in general, than the ‘groundtruth’ and SS3T CSD results, since this data-driven method discovers prior knowledge from local white matter information, i.e., white matter tracts in a compact region are likely to share a similar distribution.

We observed discrepancies between connectome reconstructed by SS3T CSD of LARDI and MSMT-CSD of HARDI, with false-positive connections (spurious connections that are not genuinely connected by fiber bundles) as well as falsenegative connections (missing connections between pairs of brain regions) when compared to multi-shell HARDI data (used here as ‘ground-truth’). These false connections were greatly reduced in connectomes based on FOD-Net of LARDI data, demonstrating the practical benefits of our proposed method. This feature is of critical importance for clinical applications of connectomics, since missing and spurious connections between different regions could inappropriately affect clinical decisionmaking.

Moreover, FOD-Net was shown to have satisfactory generalizability and reproducibility, as demonstrated in Fig. 8 using data acquired with a different scanner and protocol from the data used for training the model. Compared to SS3T CSD, FOD-Net computed more reliable connectomes regardless of the different dMRI protocols. In the test-retest experiments, FOD-Net demonstrated similar reproducibility to MSMT-CSD despite requiring substantially less data. This is because the training data used in our method includes grey matter, which can be seen as a special type of ‘data augmentation’ to perform regularization to avoid model overfitting Falk et al. (2019), and as such that it can enhance the robustness of FOD-Net. To apply our method on new subjects acquired from different scanners and clinical protocols that were not part of the training process, the steps detailed in the Section 2.3.2 (including resampling to harmonize spatial resolution) are required to obtain optimal results. As pointed out in prior literature Wang et al. (2019), using transfer learning to fine tune deep networks using samples in new datasets improves algorithm performance, and may therefore have a role to further enhance the FOD-Net model.

There are a number of potential concerns with FOD-Net. For example, a common concern for angular super-resolution algorithms in neurocomputational imaging is the potential emergence of artifacts that may not only degrade imaging quality, such as noise spikes, but also provide misleading information to clinicians (in our case, false brain connections). To explore this, we examined the macroscale structural connectomes generated from the test dMRI data. We found that only a few FOD-Net derived connections (around 4%) statistically different from the ground truth. However, in all these statistically differed regions, FOD-Net-based estimates exhibit overwhelming advantages over the state-of-the-art SS3T CSD method: even the worst results yielded by FOD-Net still outperformed SS3T CSD. Another potential concern is the formulation of FOD-Net’s angular super-resolution task, which relies on the FOD model and thus is not applicable to other dMRI models (e.g. tensor model, etc). However, FOD is now considered the de facto state-of-the-art parameterization for modeling fiber tracts Dell’Acqua and Tournier (2019) and an increasing number of clinical applications are FOD-based. Finally, our method was only trained on healthy subjects from a narrow age range (HCP young adults, aged 22-35), which may limit the application range of our model. FOD-Net might therefore not necessarily generalize to infants, older people or diseased brains, and the model may need to be trained (or transfer learning considered) for such applications.

In contrast to existing methods Lin et al. (2019); Tian et al. (2020), which strived for deep learning based dMRI angular super resolution to compensate for fiber tract quality loss induced by clinical protocols, FOD-Net directly super-resolves local fiber tracts based on the state-of-the-art mathematic model, i.e., fiber orientation distribution. This strategy overcomes the limited applicability of existing deep learning based dMRI angular super resolution methods that model raw dMRI data, which in clinical settings are inherently acquired with a diversity of protocols and a variable number of gradient directions.

Taken together, our work represents an important step to advance clinical neuroscience research by providing a deep learning - powered data enhancement framework for robust tractography and accurate connectome reconstruction. FOD-Net will improve access to high quality cerebral connectivity analysis from common clinical MRI acquisitions, potentially accelerating discovery in brain disease through the study of multicenter-multiscanner populations; and ultimately driving precision neurology through connectome analysis in routine clinical settings.

## 5. Conclusion

To the best of our knowledge, this is the first work which defines the fiber orientation distribution angular super-resolution task to generate super-resolved FOD images from low-quality single-shell LARDI FOD data, which is widely available in clinical environments. To efficiently use this novel formulation, we have proposed the novel FOD-Net, which can accurately remove spurious fibers and recovers missing fiber tracks existing in the original single-shell LARDI FOD images generated by the state-of-the-art SS3T CSD. The ease of use make FOD-Net attractive in clinical and clinical research environments for robust tractography generation and accurate connectome reconstruction. Furthermore, the pipeline of making use of FOD-Net in the clinical connectome reconstruction could serve as a useful tool for high quality cerebral connectivity analysis and brain disease detection from clinical MRI acquisitions, thus ultimately driving precision neurology through connectome analysis in routine clinical settings.

## Acknowledgments

Data were provided by the Human Connectome Project, WU-Minn Consortium (Principal Investigators: David Van Essen and Kamil Ugurbil; 1U54MH091657) funded by the 16 NIH Institutes and Centers that support the NIH Blueprint for Neuroscience Research; and by the McDonnell Center for Systems Neuroscience at Washington University, St. Louis, MO. Michael Barnett would like to acknowledge the support from the National Health and Medical Research Council of Australia & The University of Sydney Equipment Grant 2018 (grant number G201307); Chenyu Wang would like to acknowledge the support of Nerve Research Foundation, The University of Sydney, Multiple Sclerosis Research Australia and The University of Sydney – Fudan University BISA Flagship Research Program (2019); Fernando Calamante is grateful for the support of the National Health and Medical Research Council of Australia (grant numbers APP1091593 and APP1117724) and the Australian Research Council (grant number DP170101815).

